# G6PD deficiency mediated impairment of iNOS and lysosomal acidification affecting phagocytotic clearance in microglia in response to SARS-CoV-2

**DOI:** 10.1101/2023.12.12.570971

**Authors:** Abir Mondal, Subrata Munan, Isha Saxena, Soumyadeep Mukherjee, Prince Upadhyay, Nutan Gupta, Waseem Dar, Animesh Samanta, Shailja Singh, Soumya Pati

## Abstract

The glucose-6-phosphate dehydrogenase (G6PD) deficiency is X-linked and is the most common enzymatic deficiency disorder globally. It is a crucial enzyme for the pentose phosphate pathway and produces NADPH, which plays a vital role in the regulation of oxidative stress of many cell types. The deficiency of G6PD causes hemolytic anemia, diabetes, cardiovascular and neurological disorders. Notably, the patient with G6PD deficiency was severely affected by SARS-CoV-2 and showed prolonged COVID-19 symptoms, neurological impacts, and high mortality. However, the mechanism of COVID-19 severity in G6PD deficient patients is still ambiguous. Here, using a CRISPR-edited G6PD deficient human microglia cell culture model, we observed a significant reduction in NADPH and an increase in basal reactive oxygen species (ROS) in microglia. Interestingly, the deficiency of the G6PD-NAPDH axis impairs induced nitric oxide synthase (iNOS) mediated nitric oxide (NO) production which plays a fundamental role in inhibiting viral replication. Surprisingly, we also observed that the deficiency of the G6PD-NADPH axis reduced lysosomal acidification, which further abrogates the lysosomal clearance of viral particles. Thus, impairment of NO production and lysosomal acidification as well as redox dysregulation in G6PD deficient microglia altered innate immune response, promoting the severity of SARS-CoV-2 pathogenesis.

## 1. Introduction

Glucose-6-phosphate dehydrogenase (G6PD) deficiency is a common enzymopathy in the pentose phosphate pathway, affecting more than 500 million people worldwide *[1]*. The pentose phosphate pathway significantly regulates the redox microenvironment in the cells by producing reduced nicotinamide adenine dinucleotide phosphate (NADPH). Redox and inflammatory signaling pathways are essential for cellular differentiation, growth, proliferation, and immunological responses. Interestingly, NADPH shows both pro-oxidative and anti-oxidative pathways depending on the external stimuli in a tissue-specific manner *[2–4]*. Thus, NADPH’s functional dichotomy is the key to many fundamental cellular processes. However, the deficiency of the G6PD decreases cellular NADPH and dysregulates the redox microenvironment which primarily causes hemolytic anemia *[5,6]*. The World Health Organization (WHO) categorized G6PD deficiency into several classes (I to V) based on the location of mutations and degree of alteration of the enzymatic activity *[7]*. It is very evident that the degree of G6PD deficiency also represents diverse pathophysiology. Recently, diabetes, cardiovascular disorders, and neurological disorders were reported to be associated with G6PD deficiency *[8–13]*. So, G6PD-deficient patients are highly susceptible to develop diverse pathophysiology and may also show severe outcomes during infections.

Notably, multiple clinical case reports indicated high severity and mortality in SARS-CoV-2 infection among the patient having G6PD deficiency *[14–16]*. COVID-19 caused by SARS-CoV-2 affected more than 697 million people worldwide and resulted in 6.9 million deaths among the elderly and those with comorbid illnesses. Generally, SARS-CoV-2 virus enters host cells after binding to ACE2 receptors *[17,18]*. Later, the virus replicates inside the host cells which are released by exocytosis promotes pathogenesis. The SARS-CoV-2 activates innate immune responses causing activation of neutrophils and macrophages. These immune cells release proinflammatory cytokines, such as interleukins (IL-1β, IL-8 and IL-12 etc.) and TNF-α. The SARS-CoV-2-mediated inflammatory response leads to the production of reactive oxygen species (ROS) acts as a common innate defense mechanism. However, ROS are regulated by a key enzyme called G6PD via the production of NADPH, which controls the generation and removal of ROS in a tissue-specific manner. Therefore, a deficiency of G6PD can lead to the dysregulation of ROS, which causes a severe inflammatory response in COVID-19 patients. A *in-vitro* studies using lung fibroblast G6PD deficient cells showed higher infectivity of human coronavirus (HCoV) 229E *[19]*.

Notably, the G6PD deficient COVID-19 patients showed hemolysis, acute respiratory distress syndrome, prolonged ventilation support, neuropathies and high mortality *[14,20,21]*. Additionally, neurological diseases such as epileptic seizures, myelitis, cranial nerve deficit, as well as cerebrovascular disease like- stroke and transient ischemia is reported within the year after the acute SARS-CoV-2 infection in the patient with unknown G6PD genetic status *[22–25]*.

A recent report showed that SARS-CoV-2 infection provokes proinflammatory responses and apoptotic cell death in human microglia *[26]*. However, molecular mechanism of G6PD deficiency and severity of SARS-CoV-2 is not clear. Which raises question like- does oxidative dysregulation caused by G6PD deficiency only mediate severity of COVID-19 or G6PD deficiency also affects other crucial pathway important for innate immune responses. Additionally, neurological manifestation and microglial response of SARS-CoV-2 is also ambiguous in G6PD deficient condition. Therefore, we evaluated role of G6PD-NADPH axis in innate immune response in human microglia cells.

## 2. Methods

### 2.1. Culturing of Human microglia

Human microglia clone 3 (HMC3) cells from ATCC (CRL-3304) were cultured and maintained in EMEM media supplemented with 10% FBS, 1 mM Sodium Pyruvate, 1x NEAA, and 0.1% Penicillin-streptomycin. All the experiments were performed within 4 to 15 passages.

### 2.2. CRISPR-mediated G6PD deficient HMC3 line generation

Previously reported and validated sgRNA targeting exon-10 of G6PD was used to clone into the pCK002_U6-Sa-sgRNA(mod)_EFS-SaCas9-2A-Puro_WPRE vector (Addgene #85452). This particular gRNA was transfected using lipofectamine 3000 (Invitrogen #L3000001) in human microglia cells and puromycin-selected cells for 48 hours. Then cells were allowed to grow for another 10-20 days. Individual colonies were picked using local trypsinization. Further, colonies were expanded and prepared protein lysate to check G6PD level by western blot.

### 2.3. Western Blotting

Wild-type HMC3 cells and potential G6PD deficient colonies were used to make cells lysate using RIPA buffer supplemented with 1x Protease inhibitor cocktail (Roche). Protein estimation was performed using BCA method as per the manufacturing protocol (Sigma). 20µg of protein from each sample prepared in 1x protein loading dye was used for polyacrylamide gel electrophoresis (Biorad). Proteins were transferred to the PVDF membrane using a Trans-blotter (Biorad). The blot was further evaluated for G6PD expression using primary antibody against G6PD (Rabbit mAb # A11234) and the counter labelling anti-rabbit HRP secondary antibody (Invitrogen Catalog # **31460**). Chemiluminescence was capture in iBright (Invitrogen) using ECL substrate (Biorad #1705061).

### 2.4. WST-8 NADPH Estimation assay

WST-8 NADPH was performed in living cells as well in cell lysate. NADPH estimation was performed by using highly sensitive WST-8 assay and following conditions were used (Table1). Absorbance was measured at 460nm by spectrophotometer (BioTek, Synergy H1). All the experiments with biological replicates were further used to calculate SEM. Bar plots were plotted using GraphPad prism software.

### 2.5. Study the ROS by flow cytometer

1.5 x 10^5^ cells were seeded per well of 6 well tissue culture plates. On the next day, in the following condition treatments were given to cells (shown in tables). Cells were trypsinized and pelleted down by centrifugation (Eppendorf # 5810R) at 130g for 5 mins. The pellet was washed with 1x PBS two times. Finally, the pellet was dissolved in 200 µl of 1x PBS and kept in a 1.5 ml microcentrifuge tube. One of the cellular suspensions was treated with 0.1 % H_2_O_2_ for 10 minutes, this sample was used as a positive control of our experiments. Then, carboxy-H_2_DCFDA (Invitrogen #I36007) was added to the sample for labeling ROS and incubated in a 37°C incubator for 30 mins. One cellular suspension was kept as no-dye control to check the background noise. The flow cytometry was performed by Beckman Cytoflex flow cytometer.

### 2.6. In-vitro synthesis and purification SARS-CoV-2 RNA

SARS-CoV-2 Spike protein nucleotide sequence was obtained from the NCBI Gene Bank repository. It was cloned into a pSFV3 vector (Addgene #92072). SARS-CoV-2 Spike coding plasmid was received as a gift from Prof. Milan Surjit, THISTI, India. The SpeI restriction enzyme (NEB # R3133) was used to linearize the plasmid vector for *in-vitro* transcription by MEGAscript SP6 transcription kit (Invitrogen # AM1330). Then, 5’ capping of RNA was performed using the ScriptCap m^7^G Capping System (CELLSCRIPT, #C-SCCE0610). Thereafter, RNA was isolated using TRIZOL (Invitrogen #15596026).

### 2.7. Q-PCR-based gene expression study

The total RNA was isolated using TRIZOL reagent (Thermofisher part#15596206) from HMC3 cell after 24 hours treatment with various experimental conditions. The RNA concentration was measured in the Nanodrop 2000 instrument (Thermo Scientific). The RNA was used to make complementary DNA (cDNA) using a High-Capacity cDNA reverse transcription kit (Applied Biosystems # 4368814). Then, quantitative PCR (qPCR) was performed using a gene-specific primer utilizing Powerup SYBR green master mix (Applied Biosystems # A25742). Fold change (2^-ΔΔC^_t_) was calculated after normalizing with the housekeeping gene 18S in Microsoft Excel. Further, the bar plot was plotted using GraphPad Prism 9 software. The qPCR primers are mentioned in the table 3 below.

**Table1:** Experimental strategies for NADPH estimation.

| Experiments | Samples | Treatment | Duration |
| --- | --- | --- | --- |
| Validation of G6PD deficiency by measuring NADPH level | HMC3 | WST-8 +1m-PMS | 5 hours |
|  | G6PD C1 HMC3 | WST-8 +1m-PMS | 5 hours |
|  | G6PD C2 HMC3 | WST-8 +1m-PMS | 5 hours |
|  | 40µg Lysate HMC3 | WST-8 +1m-PMS | 5 hours |
|  | 40µg Lysate G6PD C1 HMC3 | WST-8 +1m-PMS | 5 hours |
|  | 40µg Lysate G6PD C2 HMC3 | WST-8 +1m-PMS | 5hours |

**Table 2:**
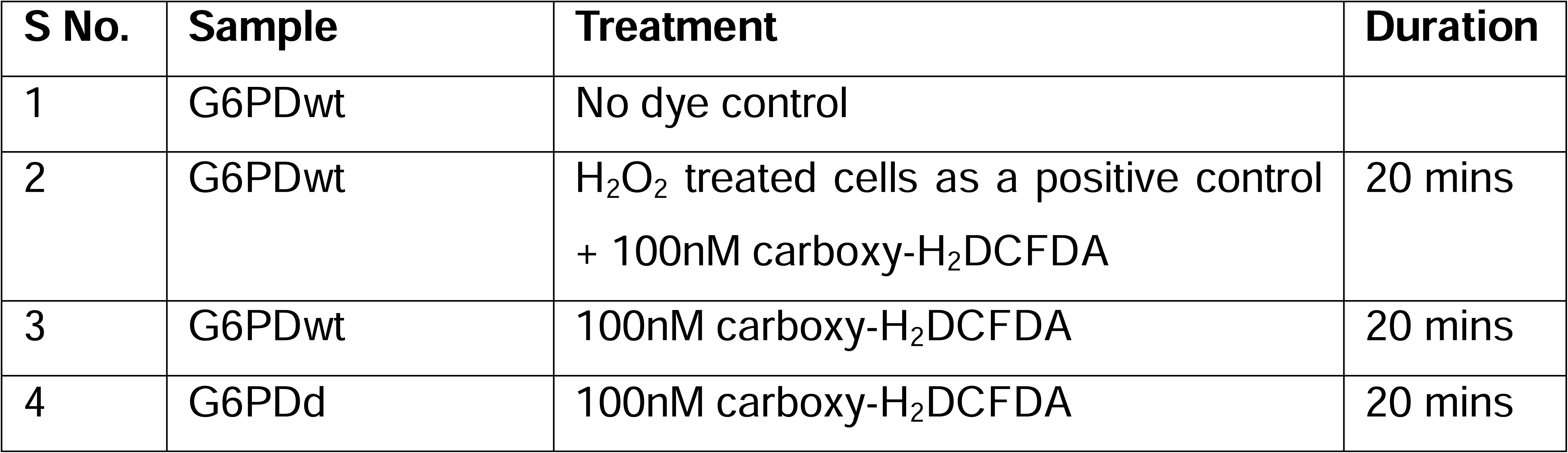
Flowcytometry strategies G6PDwt Vs G6PD deficient microglia.

**Table 3:**
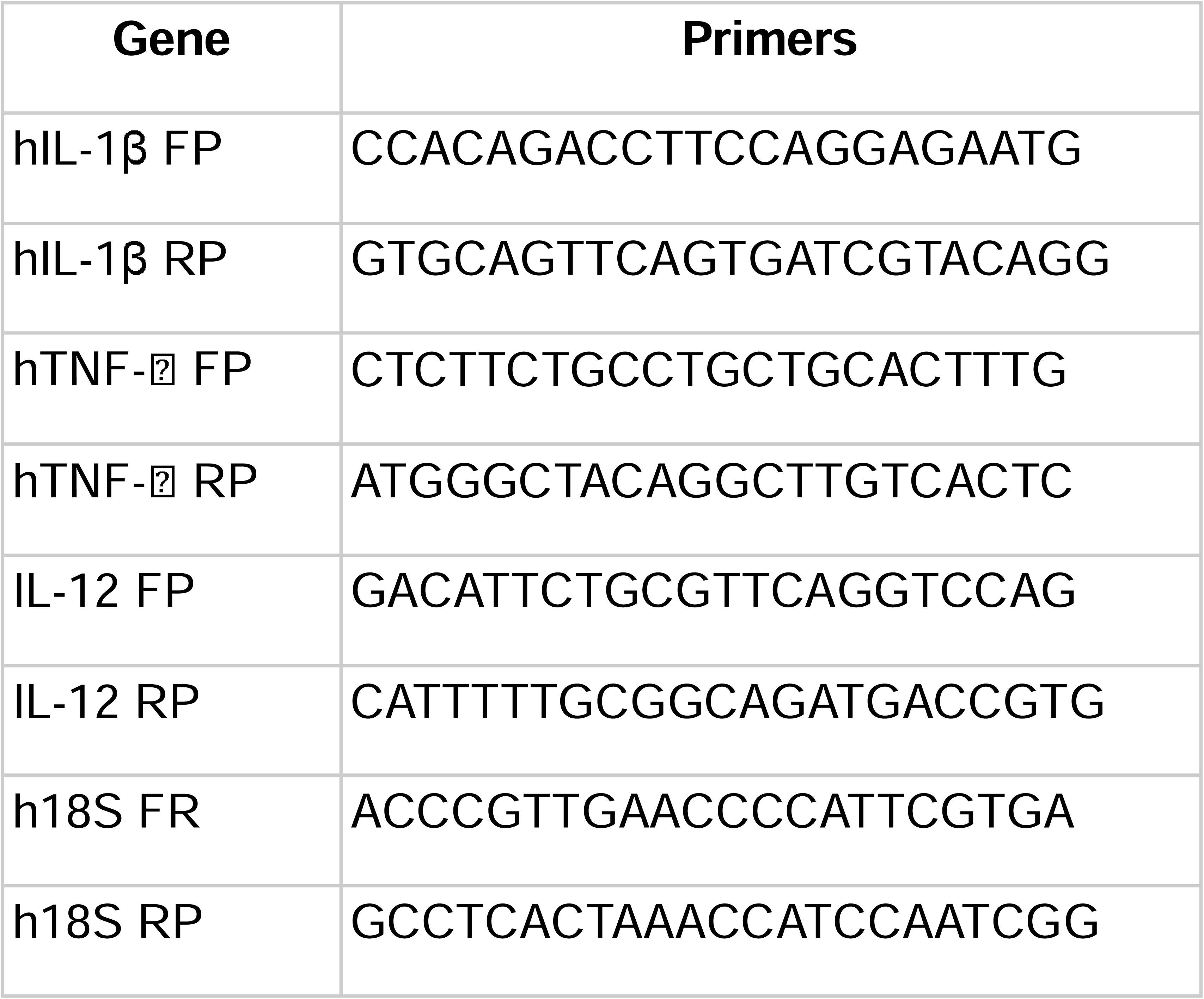
Primers details.

### 2.8. Cellular Co-localization Experiment

Confocal fluorescence intracellular Co-localization of NO with LysoTracker-deep-red: The 5 x 10^4^ HMC3 cells and cells transfectred with SARS-CoV-2 RNA were cultured on 35 mm glass-bottom Petri plates using pre-warm (37°C) complete EMEM media and the plates were incubated in 37°C CO_2_ incubator for overnight. One sets of control HMC3 cells were treated using LPS+ IFNγ (consider as a positive control). And one sets of SARS-CoV-2 transfected cells were treated with L-NAME respectively for 12 hours Then, all condtions, cells were treated with a mixture of 10 µM of NO probe and 70 nM LysoTracker (Invitrogen #L12492) prepared in cEMEM. Images were taken in the green channel (550−600 nm, ex. 488 nm) for NO and red channel (680−720 nm, ex. 640 nm) for LysoTracker deep Red (Invitrogen #L12492) by Nikon Ti2 confocal microscope.

### 2.9. Colocalization of NO with phagolysosomes using E. coli bioparticles

The 5 x 10^4^ normal HMC3 cells and HMC3 previously transfected with SARS-CoV-2 RNA were cultured on separate 35 mm glass-bottom petri plates using pre-warm (37°C) complete EMEM media and the plates were incubated in 37°C CO_2_ incubator overnight. The following day, 1:1000 ratio of pHrodo E. coli bioparticles (Invitrogen #P35361) mixture were prepared in complete EMEM media and incubated in the cells for another 12-16 hours inside the 37°C CO_2_ incubator. After washing 2-times using 1x PBS, cEMEM media containing 10 µM of NO probe along with 70 nM LysoTracker Deep Red (Invitrogen #L12492) were added to the Petri plate for 20 mins prior imaging. The images were captured in Nikon Ti2 confocal microscope.

### 2.10. Generation of mock lentivirus in laboratory

The pCK002_U6-Sa-sgRNA(mod)_EFS-SaCas9-2A-Puro_WPRE vector (Addgene #85452), pMD2G (Addgene #12259) and psPAX2 (Addgene #12260) were co-transfected in HEK293T cells using Lipofectamine-3000. After 48 hours, conditioned media of transfected cells were collected and filtered it using 0.45µ syringe filter. The viral soup was stored in - 80°C for later use.

### 2.11. Statistical analysis

All data points were collected for the calculation of standard error mean (SEM). Statistically significances were calculated based on experimental parameters by using unpaired t-test and one-way ANOVA. P-Values and statistical test were mentioned on the respective figure.

## 3. Results

### 3.1. SARS-CoV-2 spike protein expression triggers proinflammatory responses and iNOS-mediated NO production and phagolysosome formation in human microglia

Understanding the role of SARS-CoV-2 spike protein in neuroinflammation, we used a recombinant plasmid (pSFV3) carrying gene for spike protein. SARS-CoV2 spike RNA was synthesized using *in-vitro* transcription kit (Figure 1A). Later, SARS-CoV-2 RNA was transfected in the microglia using lipofectamine 2000 and the spike protein expression was confirmed by immunocytochemistry 48 hours post transfection. Our data indicated expression SARS-CoV-2 spike protein which might further cause neuroinflammation (Figure 1B). To confirm microglial activation and inflammation followed by SARS-CoV-2 RNA treatment, we performed the quantitative PCR for widely used markers such as IL1β, TNF-α and IL-12 (Figure 1C). Our qPCR data indicated increase expression of IL1β, TNF-α and IL-12 in case of SARS-CoV-2 treated microglia as compared to untreated control (Figure 1D). The LPS + IFNγ treated cells were taken as positive control for our experiments as previous reports suggested that these molecules activate microglia and promotes proinflammatory signaling. Therefore, Both SARS-CoV-2 and LPS + IFNγ showed proinflammatory response in microglia. Further, we evaluated the production of NO in microglia treated with LPS + IFNγ and SARS-CoV-2 RNA respectively. The NO production by microglia was captured by novel inhouse probe which turn on fluorescent signal after reacting with NO. The efficacy of NO detection and specificity of probe was characterized in microglia treated with LPS, IFNγ and SARS-CoV-2 RNA *[27,28]*.

**Figure 1.**
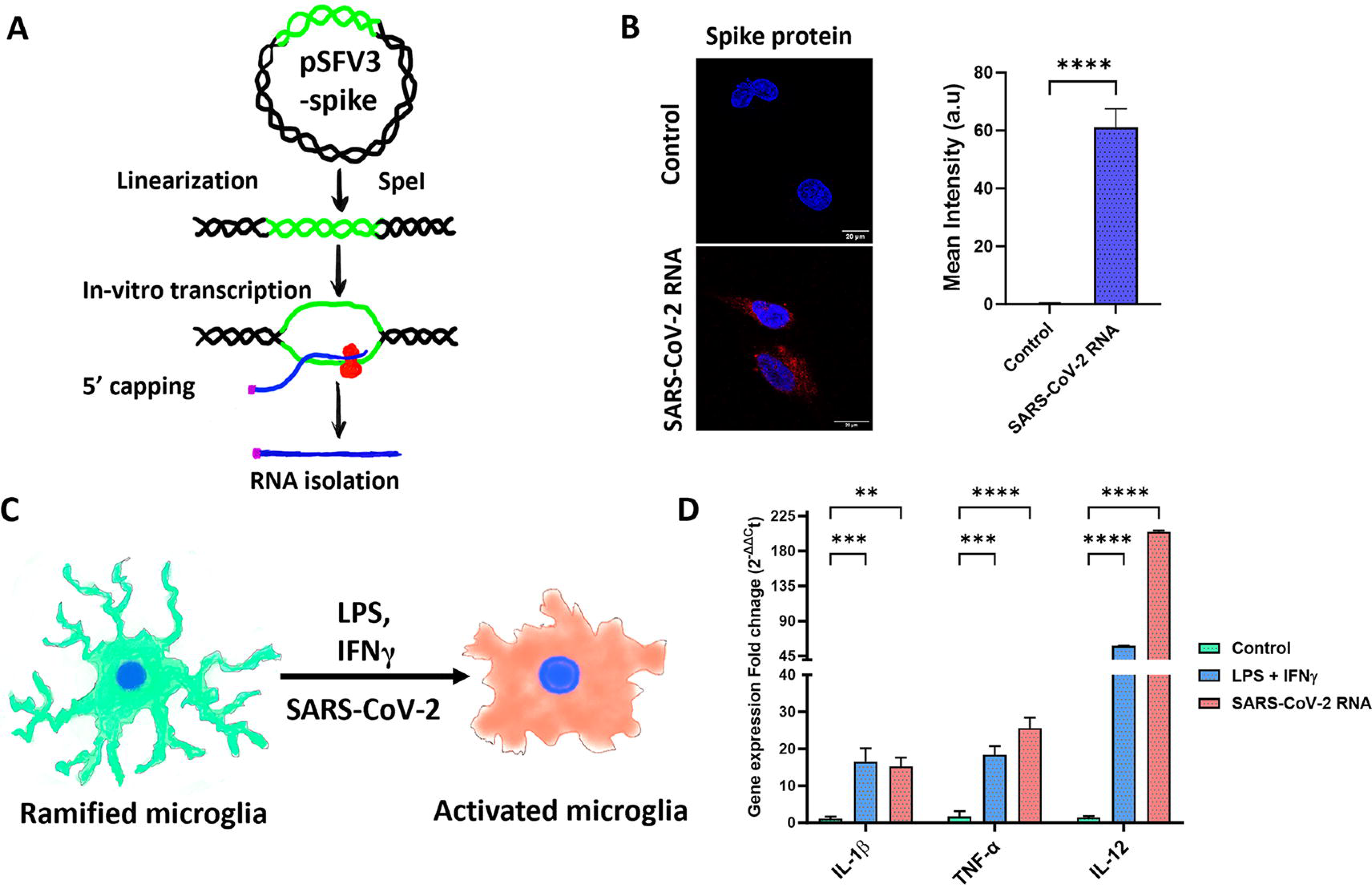
SARS-CoV-2 spike protein increases the production of proinflammatory cytokines and NO: A) A schematic representation of in-vitro transcription SARS-CoV-2 spike RNA; B) Immunofluorescence of SARS-CoV-2 Spike protein expression study by confocal microscopy after 48 hours of RNA transfection. Mean intensity was calculated from cells (n=10) Data represent the means ± SEM. The bar graph showed significance values compared to the control, with a ****p-value <0.0001 (unpaired t-test); C) Schematic representation of microglia activation using LPS, IFN-γ and SARS-CoV-2; D) Q-PCR study of proinflammatory markers expression in microglia treated with LPS + IFN-γ and SARS-CoV-2. The bar graph represents fold change in gene expression (IL-1β, IL-12, and TNF-α). Data represent the mean ± SEM (n = 3). * Represents significance values compared to control, **p<0.0016, ***p=0.005 and 0.0009, ****p<0.0001. Statistical significance was calculated using two-way ANOVA;

Moreover, we found that the microglia treated with SARS-CoV-2 spike RNA, and LPS + IFN-γ increases the production of NO as compare inactivated microglia (Figure 2A). Then, we also asked whether the generation of NO is mediated by inducible nitric oxide synthase (iNOS), as the foreign pathogens usually triggers iNOS mediated cellular host defense mechanism in majority of immune cells. The inhibition iNOS by N(gamma)-nitro-L-arginine methyl ester (L-NAME) drastically reduces NO production in microglia treated with SARS-CoV-2 RNA (Figure 2A) which suggested iNOS activation. Further, we also found the activated microglia-derived NO localized in lysosomes (Figure 2A, B). Subsequently, we evaluated the association of NO with phagolysosome of microglia using fluorescent labelled E. coli bioparticles used for phagocytosis experiment. Notably, our data indicated NO, lysosomes and E. coli bioparticles were colocalized in microglia with Pearson colocalization coefficient (r) is 0.90 in SARS-CoV-2 RNA treated cells (Figure 2C). Thus, we confirmed that NO and phagolysosome formation are highly correlated which possibly mediated clearance of foreign pathogens. However, the importance of G6PD-NADPH axis in production of NO in response to SARS-CoV-2 is yet to elucidated.

**Figure 2.**
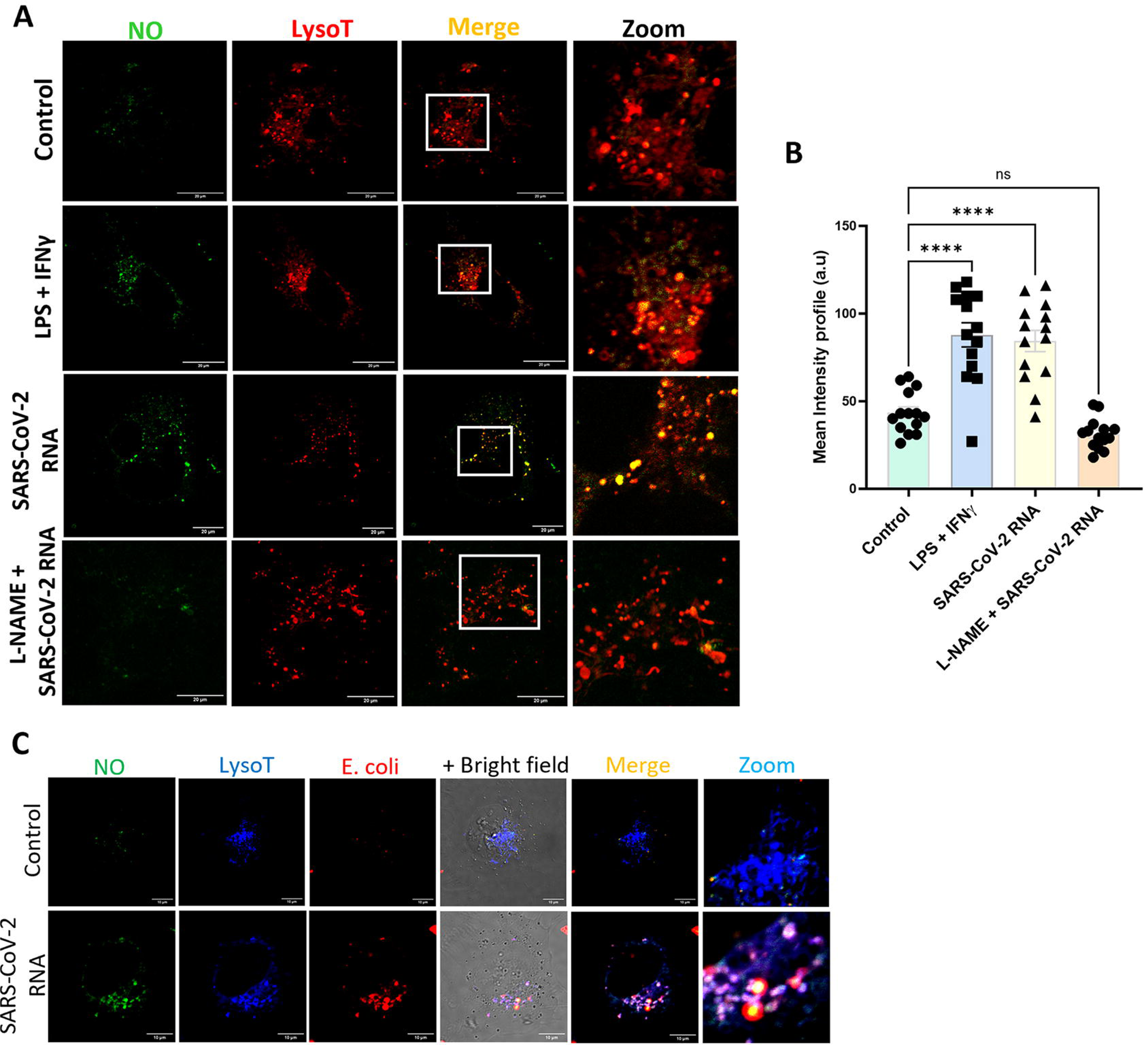
SARS-CoV-2 spike induced NO generation and sub-organellar localization in microglia. A) Confocal microscopy imaging of NO production and its colocalization with lysosomes. L-NAME was used as a known inhibitor of iNOS enzyme; B) The bar graph representing the mean fluorescent intensity of NO calculated from an equal number of cells (n=15) from each group. The statistical significance value (***P<0.001) was calculated by ordinary one-way ANOVA; C) Confocal microscopy imaging of NO, lysosomes, and E. coli bioparticle co-localization to ensure the presence of NO in the phagolysosome. The zoom image showed clear colocalization.

### 3.2. G6PD deficiency impedes NO production in microglia treated with SARS-CoV-2 spike RNA

To understand the G6PD deficiency-mediated neuroinflammation in response to SARS-CoV-2, we generated G6PD deficient human microglia cells by clustered regularly interspaced short palindromic repeats (CRISPR) tool. We used a guide RNA (gRNA) targeting exon-10 of G6PD which was previously validated by Kebin group *[29]*. We cloned the same gRNA targeting the dimerization site of G6PD exon 10 in pCK002_U6_ saCas9 puro vector. Then, microglia cells were transfected with gRNA plasmid and puromycin selection was carried out for 48 hours to pick the transfected population (Figure 3A). Afterward cells were allowed to grow for 15 days to form colonies. Multiple colonies were screened by western blotting to check the G6PD deficiency (Figure 3B). We observed a 60% reduction in the expression of G6PDd C1 and C2 as compared to control (Supplementary figure 1A). Possibly, these selected cells are producing enzymatically inactive G6PD due to Cas9 mediated *insertion-deletion* in the sequence and altered G6PD get degraded my non-sense mediated decay mechanism *[30,31]*.

**Figure 3.**
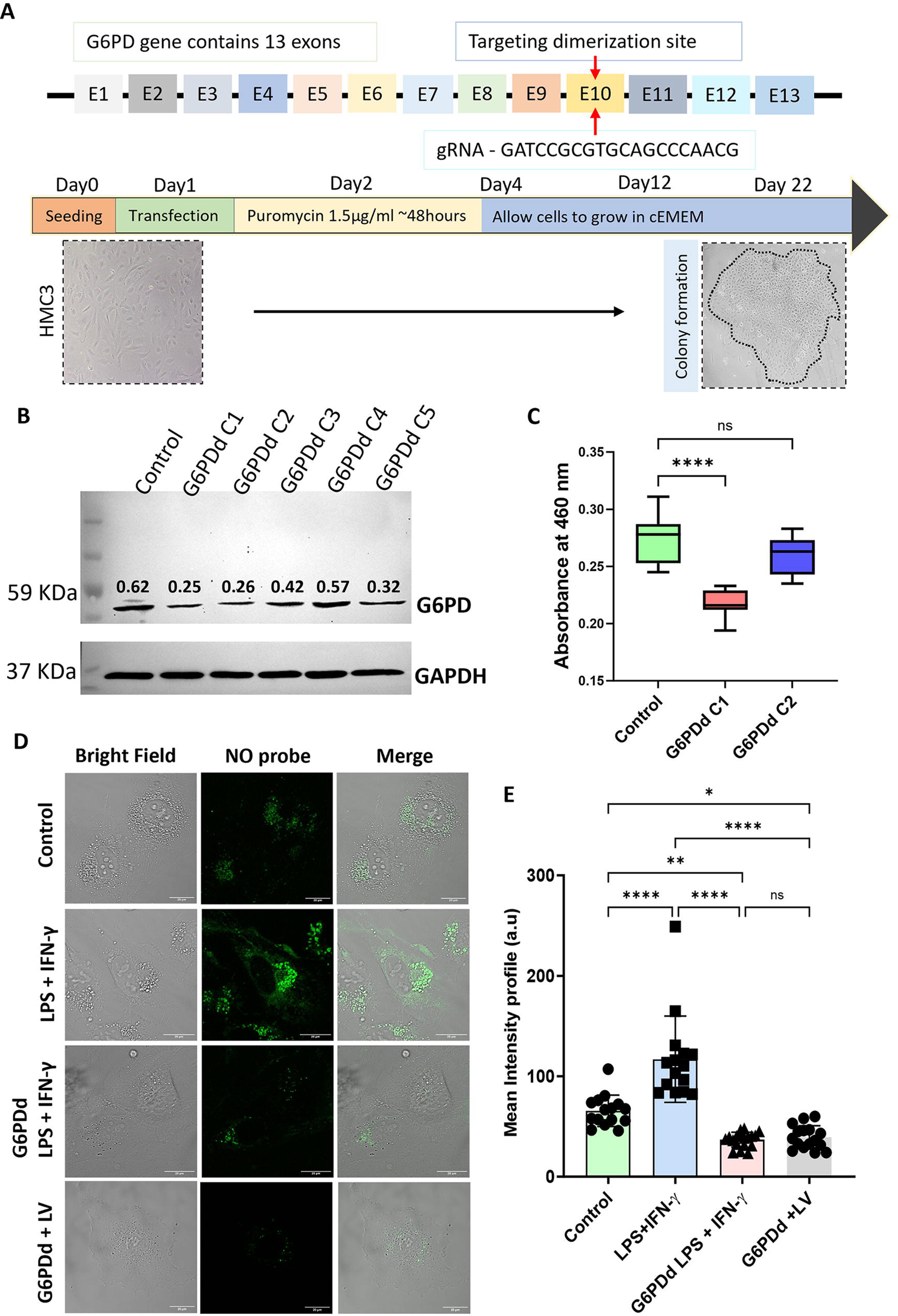
Role of G6PD-NADPH axis in NO production: A) Diagrammatic representation of G6PD genes carrying 13 exons. Previously reported gRNA was used to target the dimerization site of G6PD located on exon-10. Diagrammatic representation of CRISPR plasmid transfection followed colony expansion. B) western blot validation of G6PD deficiency in different clones; C) Box plot representing absorbance (at 460nm) value of WST-8 NADPH estimation using 40µg of cell lysate from each sample. Data represent the mean ± SEM (n= 3). *Represents significance values compared to control, ****p<0.0001. Statistical significance was calculated using ordinary one-way ANOVA; D) Confocal microscopy imaging for capturing NO production in G6PDwt and G6PDd cells under different conditions. E) The Bar graph representing mean fluorescent intensity was calculated from an equal number of cells (n=15) from each experimental group. The statistical significance value ***P<0.001, **p=0.0077, *p=0.019, ****p<0.0001 was calculated by two-way ANOVA.

Besides, we also check functional activity by measuring total NADPH production in G6PD wild type (G6PDwt) vs CRISPR edited G6PD deficient (G6PDd) microglial cells. We found a significant reduction in NADPH levels in G6PDd C1 where experiment is performed by keeping cells number equal (Figure 3C). Additionally, considering cells number may vary due to manual cell counting by hemocytometer, we also estimated NADPH in equal amount of protein from whole cell lysate which also showed significant 50 - 61% reduction in NADPH level in both G6PDd C1 and C2 (Supplementary figure 1B). Which suggested successful generation G6PD deficient microglia; However, the rest of the experiments were conducted using G6PDd C1 cells as this clone showed maximum NADPH reduction.

Moreover, we also investigated the effect of G6PD deficiency in NO production in microglia. Our data indicated, significant reduction in NO production in G6PDd as compared to G6PDwt even in presence of LPS + IFNγ and lentivirus (LV) (Figure 3D, E). Therefore, G6PD-NADPH axis is imperative for NO production in microglia. Notably, several reports showed nitric oxide inhibit viral replication and promotes clearance of viral particles *[32,33]*. So, deficiency of G6PD-NADPH-NO alters the downstream innate immune responses in microglia.

### 3.3. G6PD deficiency dysregulate oxidative stress and reduces lysosomal acidification in microglia

To probe G6PD deficiency and oxidative stress, we labelled cellular ROS by H_2_DCFDA dye and performed flowcytometry for G6PDwt and G6PDd microglia. We observed that G6PDd cells with high basal label of ROS as compared to G6PDwt microglia (Supplementary figure 1C); which suggested that G6PD deficient cells poorly regulate oxidative stress. Surprisingly, we also detected impaired lysosomal acidification in G6PDd microglia as compared to G6PDwt (Figure 4A, B). Further, exogenous supplementation of NADPH in G6PDd microglia increased lysosomal acidifications (as shown by the fluorescent intensity and bright puncta count), which suggested that the G6PD-NADPH axis plays vital role in lysosomal acidification (Figure 4A, B). However, it was unclear to us how the G6PD-NADPH axis involved in this process. From the literatures, we found a protein coded by TASL (also known as CXorf21) gene and its association with HVCN1 proton channel in the lysosomes plays vital role in lysosomal-acidification in monocyte *[34]*. Two independent studies highlighted the knockdown of TASL and HVCN1 increased in lysosomal pH in monocyte and microglia which suggested importance of these genes in lysosomal acidification *[35,36]*. Generally, TASL utilizes NADPH to generate NADP^+^, H^+^ and reactive oxygen species in the lysosomes.

**Figure 4.**
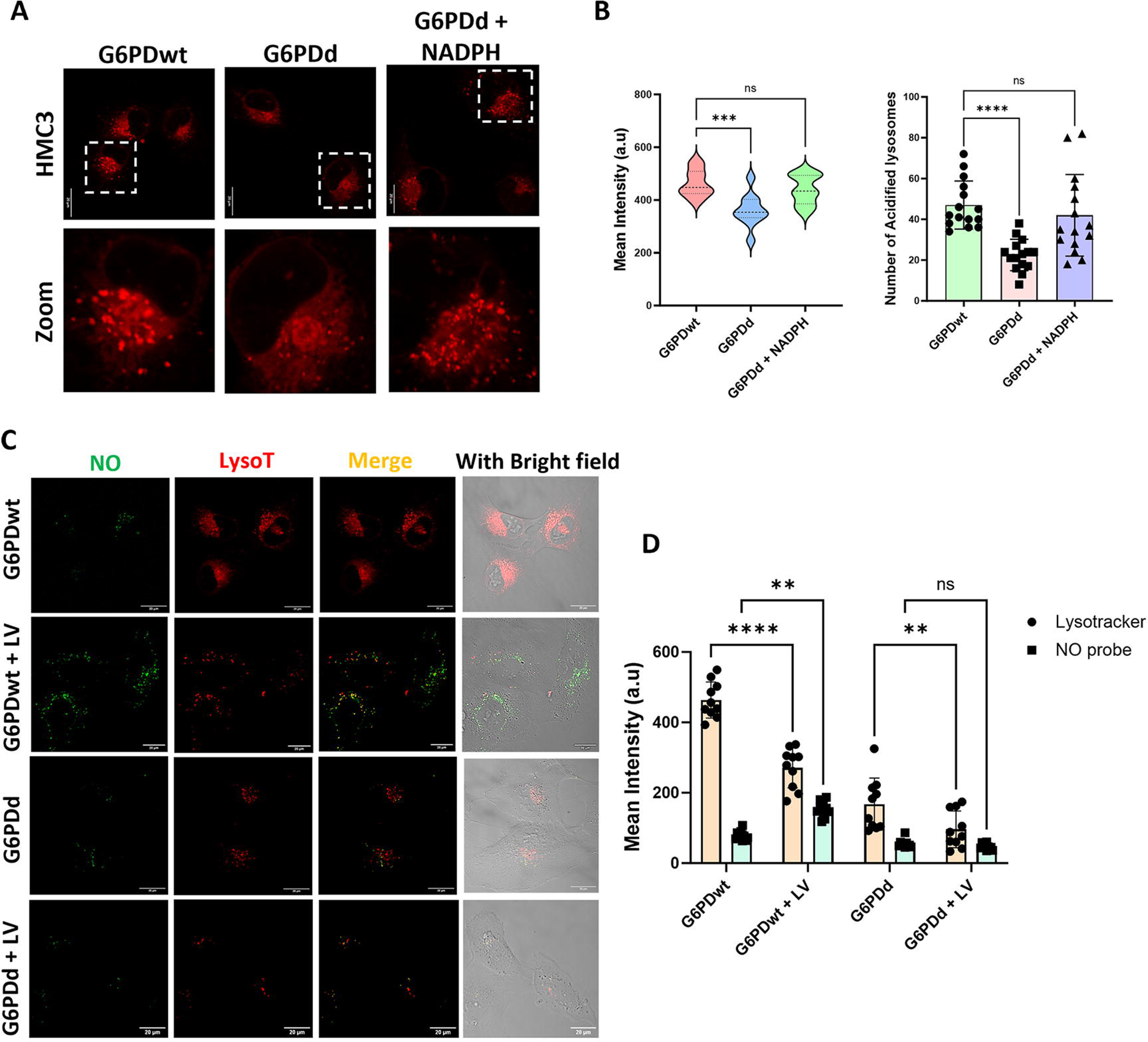
G6PD deficiency altered lysosomal function and viral clearance by microglia: A) Confocal microscopy imaging of lysosomes in G6PDwt and G6PDd and G6PDd cells supplemented with NADPH. B) The violin plot represents the mean fluorescent intensity, which was calculated from an equal number of cells (n=15) from each group. The bar graph corresponds to the number of acidified lysosomes (bright red puncta) that were also counted from the equal number of cells. The statistical significance value ***P=0.0008 and ***p<0.0001 was calculated by ordinary one-way ANOVA. C) Confocal microscopy imaging of NO and lysosome in lentivirus untreated and treated microglia in G6PDwt and G6PDd conditions. D) The mean fluorescent intensity bar plot of the NO probe and Lysotracker Deep Red was calculated from an equal number of cells (n=10). Data represent mean ± SEM. Two-way ANOVA was performed to calculate statistical significance, ****p<0.0001, **p=0.0026 and 0.0014, ns=0.63, 0.35 and 0.96.

The association of TASL with proton channel HVCN1 transports H^+^ derived from NADPH catalyzed by TASL into lysosomes and this entire process involved in lysosomal protonation/acidification and free radical generation. Therefore, the deficiency NADPH might be affecting the function of TASL and HVCN1 in microglia. However, we also analyzed the expression of TASL and HVCN1 across different brain cell types and immune cells using human protein atlas proteome database. Our analysis indicated, TASL and HVCN1 highly express in microglia compare to other brain cells (Supplementary figure 2A). And TASL and HVCN1 expression in microglia is comparable to monocyte which further suggests their key role in lysosomal function (Supplementary figure 2A, B). Therefore, the deficiency of G6PD-NADPH axis impaired the function TASL leads to decrease in lysosomal acidification in microglia. Notably, lysosomal acidification is essential for recycling of biomolecules and clearance of foreign pathogens. Thus, the deficiency of G6PD alters both NO and lysosomal function which impairs overall cellular defense machinery.

### 3.4. Viral infection in G6PD deficient microglia further abrogate phagocytotic clearance of foreign particles

Recent report shown that SARS-CoV-2 deacidified lysosome which affect the antigen presentation by host immune cells *[37]*. To study importance G6PD-NADPH axis in NO and lysosomes mediated clearance of viral particles, we used mock lentivirus prepared in our laboratory. The retrovirus such as HIV1 also deacidified lysosomes and prevent antigen processing and presentation in order to escape from immune system *[38,39]*. In our study, we observed that the mock lentivirus increases the production of NO in G6PDwt microglia. However, the mean fluorescent intensity of lysosome was reduced as compared to un-transduced G6PDwt microglia which suggested the increase in lysosomal pH in response to viruses (Figure 4C, D). Thus our results were perfectly corroborated with previous reports on impaired lysosomes acidification upon viral infection *[37,39,40]*. The addition of lentivirus on G6PD deficient cells exhibited further reduction in lysotracker mean fluorescent intensity (Figure 4C, D). Possibly, the concurrent effect of virus and G6PD deficiency completely disrupt lysosomal acidification and function in microglia. Although, we could not observe any difference in the level of NO between G6PDd and virus infected G6PDd cells; But G6PD cells infected with virus showed increased NO production. Thus, we suggest that G6PD-NADPH axis play very important role in iNOS mediated NO production and lysosomal acidification which further promotes clearance foreign particles (Figure 5).

**Figure 5.**
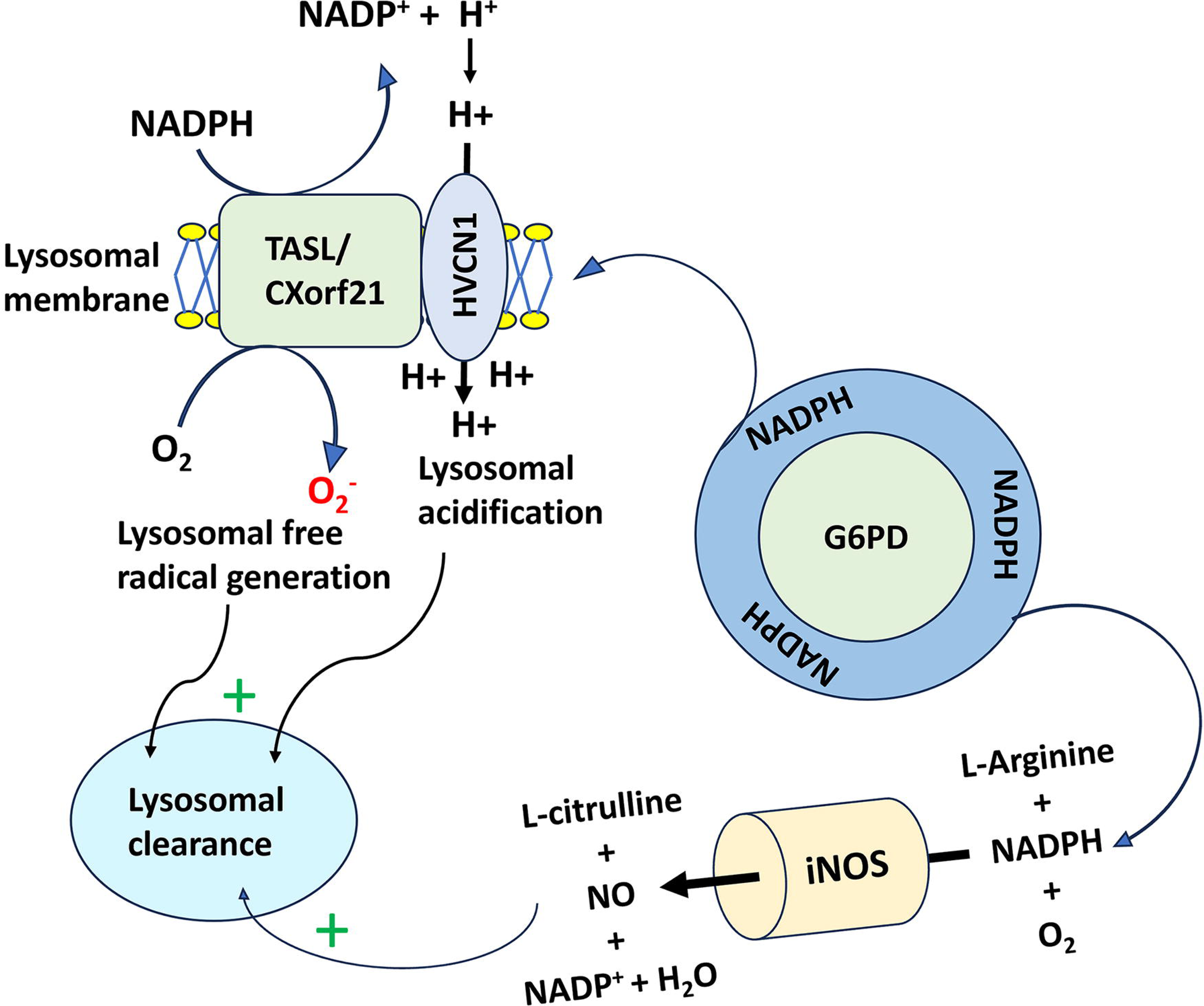
Diagrammatic representation of the role of G6PD-NADPH axis iNOS mediated NO production and TASL-HVCN1 complex mediated lysosomal acidification, which in turn help in lysosomal mediated clearance of viral particle.

## 4. Discussion

SARS-CoV-2 severity and morbidity in G6PD deficient patient raised a question on how this metabolic enzyme playing crucial role in the regulation of foreign pathogen. Our study clearly indicated that the deficiency of G6PD decrease cellular NADPH level and increase basal ROS which further suggested poor redox management in microglia. Besides, NO is crucial for the regulation of viral pathogenesis, and which is significantly downregulated in G6PD deficient condition. So, exogenous supplementation of NO might be very helpful in G6PD deficient patient during any pathogen’s infection. Additionally, we also observed the lysosomal deacidification in G6PD deficient microglia which added another crucial function of G6PD-NADPH axis in cells. Notably, various report showed that several virus including SARS-CoV-2 also causes deacidification of lysosomes and followed egress pathway *[40,41]*. A recent report suggested that a distinct motif in the E-protein of SARS-CoV-2 is required for viral particle formation and lysosomal deacidification in host cells *[40]*. Thus, entry of virus and its replication dysregulate lysosomal function and antigen presentation process by host immune cells. However, apart from virus mediated lysosomal deacidification, we also found that the G6PD-NADPH axis play substantial role in lysosomal acidification. Thus, G6PD deficiency can be considered as a genetic co-morbid factor for any pathogen invasion into body. Several antioxidative therapeutics such as N-acetyl cysteine (NAC), tocopherol was used for the treatment of G6PD deficiency which did not show any promising result *[42]*. Therefore, the unavailability of therapeutics for G6PD deficiency made more challenging. Future research should focus on development of G6PD therapeutics that will potentially allow us to prevent the diverse pathogenesis of G6PD deficiency. However, NO inhalation showed effective therapeutics against COVID-19 irrespective of patient’s genetic background which could also be very beneficial for G6PD deficient patients *[43]*. Additionally, AG1, a small molecule activator increase the enzyme activity of G6PD clinical variant which can be useful for restoration of NADPH level in cells *[44]*.

## Acknowledgment

Ph.D. students A.M, SM, SM and WD are grateful to Shiv Nadar Institute of Eminence Deemed to be University (SNIoE), Delhi NCR, and the Shiv Nadar Foundation for their research fellowships. PhD student PU is grateful to University Grant commission (UGC) for fellowship. We acknowledge Prof. Milan Surjit for providing a pSFV2 vector containing the gene for the SARS-CoV-2 spike protein as a gift. The authors acknowledge the DST-FIST grant [SR/FST/LS-1/2017/59(c)] for the confocal microscopy facility at Shiv Nadar Institute of Eminence Deemed to be University, Delhi NCR. We also acknowledge Shiv Nadar Foundation Core Research grant.

## Authors contribution

Conceptualization: AM, SS and SP; Investigation and methodology: AM, SM, IS, SM, PU, NG, WD; Data analysis and Manuscript writing: AM, Manuscript Editing: SS, SP; Funding acquisition and supervision: SS, SP, AS; All authors have read and agreed to the final version of the manuscript.

---

**Supplementary figure 1.**
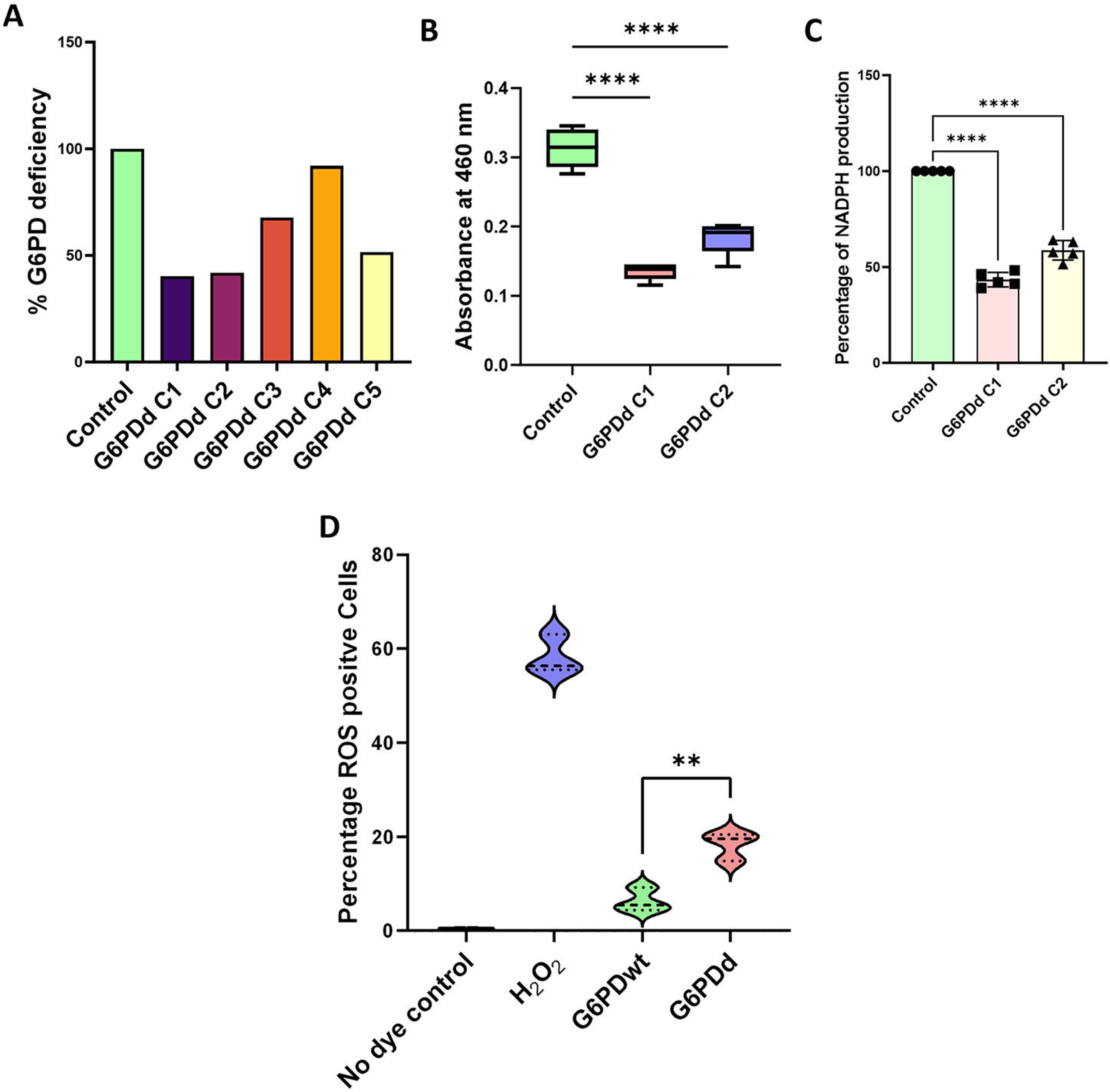
A) Bar graph represents the percentage of G6PD deficiency calculated from integrated densitometry value from western blotting; B) Box plot representing WST-8 NADPH estimation using an equal number of cells (4 x 10 HMC3 cells) in each well; C) Bar graph representing the percentage of NADPH level in G6PDwt Vs G6PDd cells. Data represent mean ± SEM. Ordinary one-way ANOVA was performed to calculate significance value (****p<0.0001); D) Violin plot representing the percentage of ROS-positive cells in each experimental condition. Data represent mean ± SEM. Statistical significance was calculated using an unpaired t-test, **p=0063.

**Supplementary figure 2.**
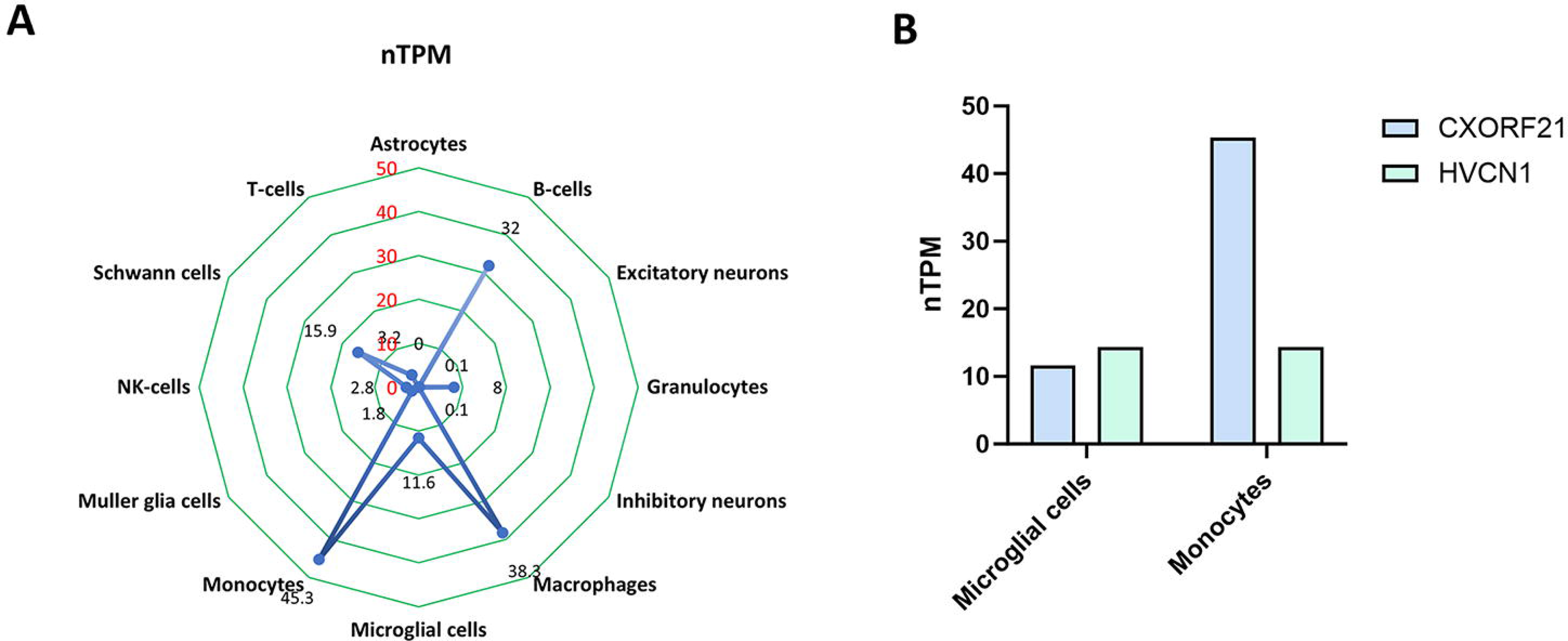
A) The radar plot representing the expression of TASL/CXorf21 from different brain and immune cell types generated from the normalized transcript per million (nTPM) gene expression values from the Human Protein Atlas Database. B) The bar diagram represents the expression (nTPM expression value from the Human Protein Atlas database) of TASL/CXorf21 and HVCN1 in monocyte vs microglia.

